# High resolution parallel sequencing reveals multi-strain *Campylobacter* in broiler chicken flocks testing ‘negative’ by conventional culture methods: implications for control of *Campylobacter* infection

**DOI:** 10.1101/2022.03.28.485779

**Authors:** Frances M. Colles, Daniela Karasova, Magdalena Crhanova, Stephen G. Preston, Adrian L. Smith, Marian S. Dawkins, Ivan Rychlik, Sabine G. Gebhardt-Henrich

## Abstract

Contaminated chicken meat is a major source of human Campylobacteriosis and rates of infection remain high, despite efforts to limit the colonisation of broiler (meat) chicken flocks on farms. Using conventional testing methods of culture or qPCR, *Campylobacter* is typically detected amongst broiler flocks from 3 weeks of age, leading to the assumption that infection is introduced horizontally into chicken rearing houses at this time. In this study, we use parallel sequencing of a fragment of the *Campylobacter* outer membrane protein, encoded by the *porA* gene, to test for presence of *Campylobacter* DNA amongst fresh faecal samples collected from broiler flocks aged 23-28 days. *Campylobacter* DNA was detected in all of the 290 samples tested using the *porA* target, and in 48% of samples using 16S bacterial profiling, irrespective of whether or not *Campylobacter* could be detected using conventional qPCR thresholds. A single *porAf2* variant was predominant amongst flocks that would be determined to be *Campylobacter ‘*positive’ by conventional means, but a diverse pattern was seen amongst flocks that were *Campylobacter* ‘negative’. The ability to routinely detect low levels of *Campylobacter* amongst broiler flocks at a much earlier age than would conventionally be identified requires a re-examination of how and when biosecurity measures are best applied for live birds. In addition, it may be useful to investigate why single *Campylobacter* variants proliferate in some broiler flocks and not others.

## INTRODUCTION

Raw or uncooked poultry meat has been identified as the main route by which humans become infected with *Campylobacter*, one of the major causes of gastroenteritis (Sheppard, et al., 2009). In the European Union, over 70% of broiler (meat) chicken flocks were found to be *Campylobacter* positive at the time of slaughter (European Food Safety Authority, 2010). The EU prohibits disinfection of chicken carcases by chlorine which means that contaminated chicken can easily make its way onto supermarket shelves. Public Health England (now the UK Health Security Agency), for example, found *Campylobacter* in 73% of supermarket chicken and on 7% of the outer packaging (Jorgensen, et al., 2019).

Using standard culture or qPCR methods, *Campylobacter* are not usually detected in chicken flocks until the birds are at least 3 weeks of age. We henceforth refer to *Campylobacter* status based upon these methods as *‘Campylobacter* culture/qPCR positive/negative’ for clarity. The widespread use of such methods as the main way of detecting the presence of *Campylobacter* gave rise to the assumption that because the bacteria could not be detected until this age, they were not present in the birds. It therefore seemed logical that the key to controlling *Campylobacter* was to prevent infection from outside sources by stricter biosecurity. However, the combination of continuing *Campylobacter* infection despite the introduction of tighter biosecurity measures (Anonymous, 2017) and sensitive genetic techniques for detecting the bacteria has challenged the idea of initially pristine flocks becoming later contaminated from outside due to a breach in biosecurity (Colles, et al., 2021; Cox, et al., 2012). Using a deep sequencing approach, Colles *et al*. (2021) detected *Campylobacter* DNA in faecal samples from all the broiler flocks they tested when birds were less than 8 days old (Colles, et al., 2021). *Campylobacter* DNA was detected amongst a total of 87.5% of 16 broiler flocks from the UK, Switzerland and France by 16S bacterial profiling assay and amongst 100% of 34 flocks using the *porAf2* assay. The amount of *Campylobacter* DNA identified in these young flocks was very small (typically less than 0.01% of the microbiome) and it was also notably diverse, showing wide variation at the *porA* locus. By contrast, flocks that later (at 28-46 days old) tested positive for *Campylobacter* using culture or PCR assay had an average 100 fold increase in *Campylobacter* DNA (perhaps associated with ‘super-shedder’ individuals), but predominantly of one *porA* genotype. Furthermore, >28 day old flocks that tested negative using culture or qPCR tests retained the juvenile pattern of very small quantities of *Campylobacter* DNA of high genetic diversity.

These findings suggest new strategies for the control of *Campylobacter*. If *Campylobacter* are universally present in chicken flocks by the time the birds are a week old (and possibly earlier), but not all flocks later develop infections severe enough to be detected as ‘positive’ by culture or qPCR, then it might pay to investigate why some flocks retain the early pattern of low levels of diverse *Campylobacter* and others develop high levels of a single strain. As higher levels of *Campylobacter* are likely to be most dangerous to humans through increased risk of transmission, understanding what triggers the increase could be the first step in reducing the risk of human infection.

In this paper, we provide further supporting evidence that at 28 days of age, flocks testing positive for *Campylobacter* via culture or qPCR tests are shedding large quantities of mainly single strains of *Campylobacter*, although precisely which strain they shed varies from flock to flock. At the same time, flocks testing negative by the same conventional tests are also shedding *Campylobacter* as detectable by deep sequencing but only in minute quantities and of diverse strains. We also show that different testing methodology for *Campylobacter* can give different results as to whether a flock is classified as positive or negative and argue that the resulting uncertainty over the *Campylobacter* status of a flock may be one reason why the search for effective control measures has proved so difficult.

## MATERIALS AND METHODS

### Ethical approval

The study was approved by the Institutional Review Board for the approval of animal experiments of the LANAT office of the Canton of Bern (BE97/16) and met all cantonal and federal regulations for the ethical treatment of animals on 30-09-2016. The procedure was declared to be severity level 0.

### Sample collection

Fresh faecal samples were collected from 20 flocks from three different farms; 8 from Farm 1, 7 from Farm 2 and 5 from Farm 3. Details of housing and management were described by Dawkins et al. (2021). Samples were collected between November 2017 and October 2018, with 5 flocks sampled in Winter months (December, January, February), 4 flocks sampled in Spring months (March, April, May), 4 flocks sampled in Summer months (June, July) and 7 flocks sampled in Autumn (September, October, November). Up to 16 samples were collected per flock, when the birds were aged between 23 and 28 days of age (Table S1, supplementary data). Samples were stored at −20°C in RNA*later*® before shipping on dry ice for DNA extraction.

### DNA extraction

Caecal content samples were homogenised in a MagNALyzer (Roche, Basel, Switzerland). Following homogenisation, the DNA was extracted using a QIAamp DNA Stool Mini Kit according to the manufacturer’s instructions (Qiagen, Hilden, Germany). The DNA concentration was determined spectrophotometrically and DNA samples diluted to 5 ng/ml.

### qPCR

DNA extracts from samples were tested individually using the method published previously (Colles, et al., 2021). Presence of *C. jejuni* DNA was detected using primers/probe for the *mapA* gene and the presence of *C. coli* DNA detected using primers/probe for the *ceuE* gene (Best, et al., 2003). Positive results were recorded for Ct values of 16 to 35, corresponding to copy number >100, based upon plasmid controls. All results, including detection of lower copy number, with higher Ct values are given in Table S1, supplementary data.

### 16S bacterial profile sequencing

The V3/V4 region of 16S rRNA genes were amplified using the method described previously (Kubasova, et al., 2019). Briefly, the following primers were used, with MIDs representing different 5, 6, 7 or 9 base pair sequences to allow multiplexing of samples; forward primer 5’-TCGTCGGCAGCGTCAGATGTGTATAAGAGACAG-MID-GT-CCTACGGGNGGCWGCAG-3’ and reverse primer 5’-GTCTCGTGGGCTCGGAGATGTGTATAAGAGACAG-MID-GT GACTACHVGGGTATCTAATCC-3’. The KAPA HiFi Hot Start Ready Mix kit (Kapa Biosystems, Woburn, MA, USA) was used for PCR amplification, and the resulting PCR products were purified using AMPure beads. The PCR products were indexed using the Nextera XT Index Kit following the manufacturer’s instructions (Illumina, San Diego, CA, USA) and the concentration of the differently indexed samples determined using a KAPA Library Quantification Complete kit (Kapa Biosystems, Woburn, MA, USA). Sequencing was performed using the MiSeq Reagent Kit v3 (600 cycle), with 20 pM phiX DNA added to the pooled indexed PCR products to give a final concentration of 5% (v/v). Quality trimming of the raw reads was performed using TrimmomaticPE v0.32 with sliding window 4 bp and quality read score equal or higher than 20 (Bolger, et al., 2014). Minimal read length was at least 150 bp. The fastq files generated after quality trimming were uploaded into QIIME software (Caporaso, et al., 2010). Forward and reverse sequences were joined and in the next step, chimeric sequences were predicted and excluded by the slayer algorithm. The resulting sequences were then classified by RDP Seqmatch with an OTU (operational taxonomic units) discrimination level set to 97 %.

### Campylobacter porA parallel sequencing

A short fragment of the short variable region of the *porA* gene (“*porAf2”)* was amplified in triplicate 25µl reactions, using the method published previously (Colles, et al., 2019). The fragment is specific for *C. jejuni* and *C. coli;* no-cross reaction to other bacterial species was detected by BLAST searching the NCBI nucleotide database, or amongst whole genome sequence from a further 40 *Campylobacter* species, *Helicobacter pullorum or H. pylori*, tested using the multispecies Ribosomal MLST database (Jolley, et al., 2012). Primers were designed to match short sections of *porA* nucleotide sequence data without polymorphisms, covering both *C. jejuni* and *C. coli*. Briefly, the MOMP B 5’-CCA CAA TTA TGG TTA GCT TA −3’ and MOMP 2R 5’-TGA GAA GTT AAG TTT TGG AGA G-3’ primers were used, with the MOMP 2R primer tagged with a 7 nucleotide barcode specific for each reaction, enabling the reactions to be multiplexed within the same sequencing library. The PCR mastermix was made according to the manufacturer’s recommendations, using high fidelity Phusion Hot Start Flex DNA polymerase enzyme and 5X Phusion HF buffer (New England Biolabs, UK, M0535). Library preparation was performed using the NEB ultra DNA library preparation kit for Illmina (E7370) and Indexing primers (E7335S and E7500S) (New England Biolabs Inc, UK), following protocols described previously (Colles, et al., 2019).

PCR products were loaded with 10% phiX onto the Illumina MiSeq platform, following analysis by TapeStation (Agilent Genomics) and qPCR (E7630C, NEB) to confirm template size and concentration. The 600-cycle MiSeq Reagent Kit v3 (Illumina, UK, MS-102-3003) was used, giving paired 300 nucleotide reads. Raw sequencing reads were processed using the DADA2 pipeline (Callahan, et al., 2016) and then the phyloseq package in R was used to produce a table of OTUs (McMurdie and Holmes, 2013), as published perviously. Custom python scripts were used to demultiplex reads according to their barcode, and cutadapt v1.15 (Martin, 2011) was used to trim any remaining primer sequence ahead of the DADA2 pipeline. OTU’s were assigned both *porAf2* nucleotide allele and its translated MOMPf2 peptide allele using the PubMLST database (https://pubmlst.org/organisms/campylobacter-jejunicoli/). The assigned alleles and sequence information are publicly available on the database by searching ‘Typing’-’Downloads’-’Loci not in schemes’.

### Cross-contamination control

The parallel sequencing for *Campylobacter porA* was performed in a separate institute to the bacterial 16S profiling, but using aliquots of the same DNA extractions. PCR reactions were performed in separate PCR cabinets for mastermix preparation and template addition, within a designated clean room. Equipment was cleaned before use and between batches of samples using DNA*Zap*^*TM*^ (AM98902, ThermoFisher Scientific, UK), 70% ethanol and UV light for a minimum of 15 minutes. Fresh aliquots of reagents and newly opened plastic ware were used for each set of PCR reactions, and non-template controls with molecular water used in place of sample were included for every batch. Sequencing reactions were prepared in a separate room to that of the PCR reactions.

### Data analyses

Data were transformed to an even sampling depth, giving proportional frequency of each *porAf2* type, using the phyloseq package in R (McMurdie and Holmes, 2013). Excel and Tableau 2019.4 software were used to produce colour matched bar charts. The Simpson’s and Shannon’s diversity indices were performed on raw and interpolated/extrapolated data calculated the iNEXT (iNterpolation/EXTrapolation) R package to ensure standardised comparison (Hsieh, et al., 2016). For Simpson’s diversity index, a 1-*D* value of 1.0 indicated that all members of a population could be distinguished from each other, and a 1-*D* value of 0 indicated that all members of a population were identical (Hunter, 1990). Shannon’s diversity index was included as it is considered to give more weight to rare species (*pofAf2* variants) (Shannon, 1948). An H value of 0 indicated that all species were the same. H increases with increasing number of species. Spearman’s rank correlation coefficient measuring the relationship between qPCR value and % *Campylobacter* 16S DNA was calculated using R. Bray-Curtis dissimilarity indices were calculated using the Phyloseq package in R (McMurdie and Holmes, 2013) for the *porAf2* populations identified in each flock, and compared by flock, ‘florid/non-florid’ status (predominance or otherwise of a single *porAf2* variant), and parent flock. The results were plotted using ggplot2 (Wickham, 2016) as non-metric multidimensional scaling (NMDS) ordination plots. The evolutionary distances between the *porAf2* variants were calculated using the Neighbour joining and p distance methods (Saitou and Nei, 1987), using the MEGA-X software (Kumar, et al., 2018).

## RESULTS

### Detection of Campylobacter DNA by qPCR, parallel sequencing of the 16S ribosomal RNA and porAf2 gene fragment targets

Using Ct value thresholds of 16-35, corresponding to copy number >100, > 90% of samples from three broiler flocks (flocks 3, 14 and 19) were determined to be positive for *Campylobacter*, using the *mapA* and *ceuE* targets. Two other flocks recorded Ct values of 31-34, corresponding to ∼1,000-30,000 copy number for 4/16 (25%) samples (flock 7) and 6/16 (37.5%) samples (flock 11). Samples were collected from these flocks in June (farm 1), May and September 2018 (farm 2) and February and December 2018 (farm 3). If the Ct threshold was removed, and less stringent accuracy therefore adopted, *Campylobacter* DNA was detected amongst at least one sample from all of the 20 flocks tested at 23-28 days of age by qPCR. Of these, 14 (70%) flocks were positive for both *C. jejuni* and *C. coli*, 4 (20%) flocks were positive for *C. jejuni* only and 2 (10%) flocks were positive for *C. coli* only. *Campylobacter* DNA was detected amongst 145 of the 290 (50%) samples tested individually by qPCR. Of these, both *C. jejuni* and *C. coli* were detected amongst 39 (26.9%) of 145 samples, *C. jejuni* only was detected amongst 72 (49.6%) of 145 samples and *C. coli* only was detected amongst 35 (24.1%) of 145 samples (Table S1, supplementary data).

Variants of *Campylobacter* 16S rDNA were recovered from at least one sample from 17 of the 20 (85%) flocks, and 142 of the 291 (48.8%) samples tested. Amongst the positive samples, the proportion of *Campylobacter* DNA identified amongst the bacterial 16S variant sequences ranged from < 0.01 to 79.03% per sample (Table S1, supplementary data). The number of ‘positive’ samples/birds ranged from one bird within a flock, to all 16 birds tested from a flock. Presence/absence of *Campylobacter* DNA matched by 16S bacterial variant profile and low stringency qPCR in 176 (60.7%) of the 290 samples. *Campylobacter* DNA was detected in 43 (14.8%) of 290 samples by low stringency qPCR but not by 16S bacterial profile, and in 72 (24.8%) of 290 samples by 16S bacterial variant profile but not by qPCR. The different methods of *C. jejuni* and *C. coli* quantification by qPCR and the percent *Campylobacter* DNA detected by 16S rDNA profile were correlated (when comparing 16S rDNA with qPCR results for *C. jejuni* alone (Spearman’s rank correlation coefficient *ρ* =0.47, *p* <0.01), and qPCR results for *C. jejuni* and *C. coli* combined (*ρ* =0.38, *p* <0.01)) amongst the samples tested.

*PorAf2* variants were detected from all of the samples from all of the flocks tested in the study, with an average sequencing depth of 2,000 (range 31-220,000) per sample.

### Diversity of porAf2 variants

A total of 245 *porAf2* nucleotide variants were recovered from the samples and included in the study (Figure 1). Of these, 149/245 (60.8%) *porAf2* variants were newly described in this study compared to previous. The number of different *porAf2* variants recovered from an individual sample ranged from 1 to 79, with an average of 21.1. The *porAf2* variant types 1-9 were most commonly isolated in the study, accounting for 70.5% of the total sequences recovered (Figure 2). The 245 *porAf2* variants translated to 170 peptide variants, with the 79 *porAf2* variants amongst an individual bird translating to 65 different peptide sequences. Point mutations and deletions were spread along the *porAf2* fragment sequenced, with the most disparate alleles varying by ∼80% sequence homology.

**Figure 1.**
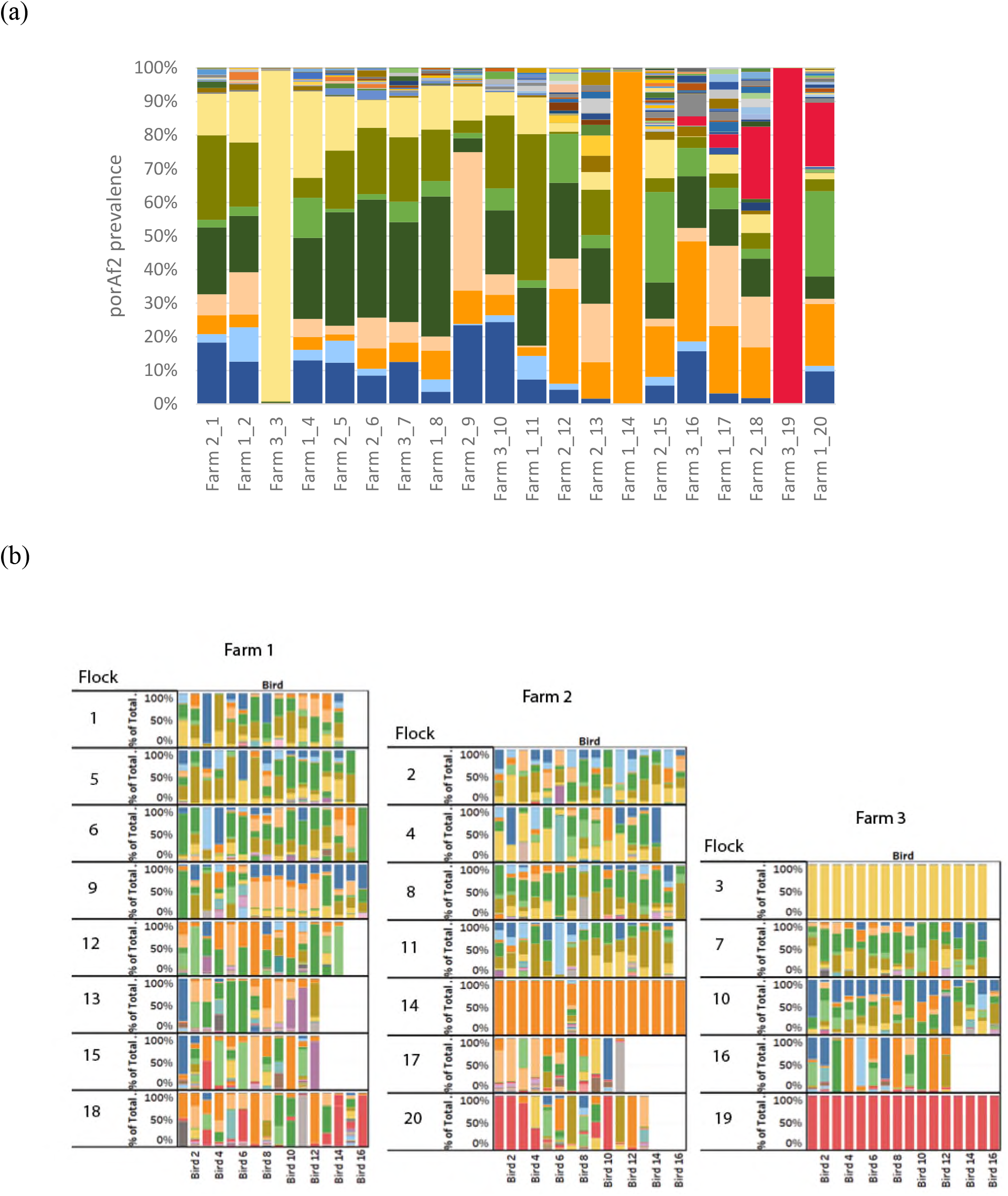
The *Campylobacter porAf2* nucleotide variants shown by (a) Farm_flock id, and (b) individual sample, with each colour representing a different variant.

**Figure 2.**
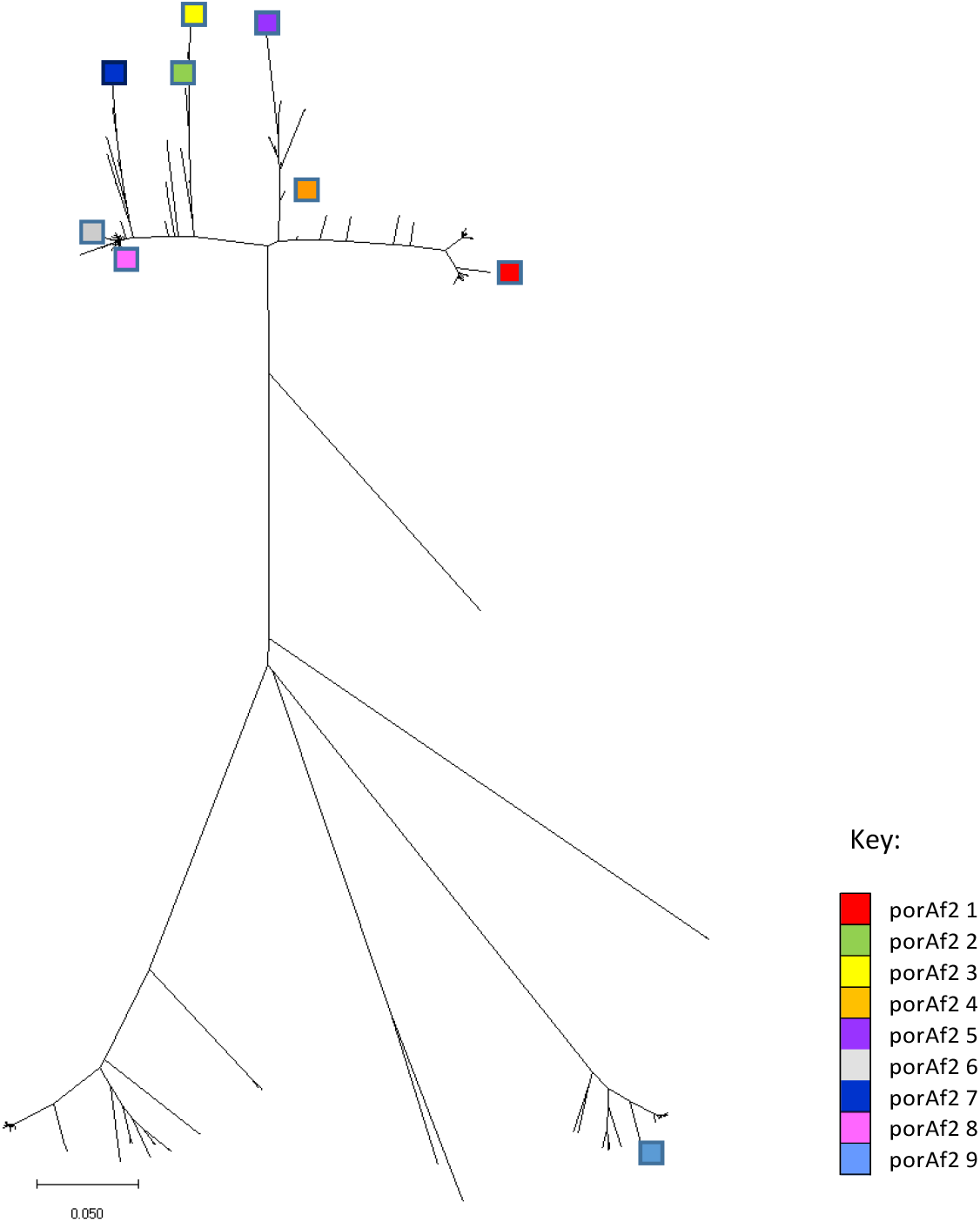
A neighbour joining tree showing the genetic distance between the 245 *porAf2* variants identified amongst the broiler flocks. The most commonly isolated *porAf2* variants 1-9 are highlighted in colour.

In three flocks (flocks 3, 14 and 19), a single *porAf2* variant was predominant amongst all of the samples tested (Figures 1 and 2). These three flocks also had the highest levels of *Campylobacter* detected by qPCR, percentage of *Campylobacter* amongst 16S bacterial variant sequence and number of birds testing positive within a flock. *Campylobacter* 16S DNA was not detected in flock 7 that had 4 positive samples by qPCR, and at very low quantities <1% in flock 11 that had 6 positive samples by qPCR. The *porAf2* variants were more evenly spread amongst samples for each of the remaining samples and flocks, excluding those from flocks 3, 14 and 19. The Simpson’s and Shannon’s diversity indices were close to zero for all except one bird/sample in flocks 3, 14 and 19 (Figure 3). For these three flocks, the Simpson’s diversity ranged from <0.01-0.04 (0.81 for the outlying sample), with an average of 0.03, and the Shannon’s diversity ranged from 0.03-0.15 (2.28 for the outlying sample), with an average of 0.12. For the remaining flocks, the Simpson’s diversity ranged from 0-0.91, with an average of 0.59, and the Shannon’s diversity ranged from 0-2.57, with an average of 1.22.

**Figure 3.**
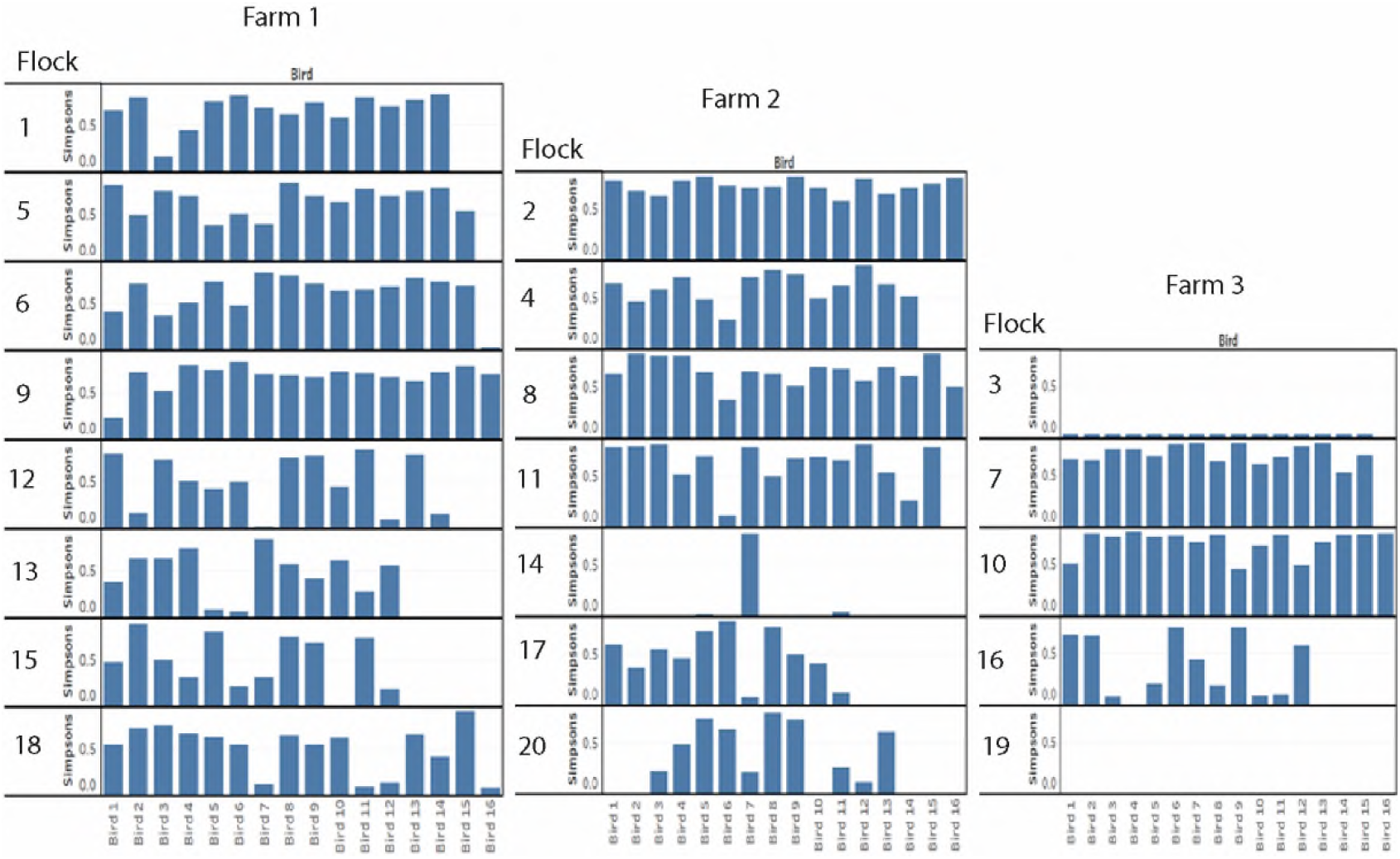
Measures of Simpson’s diversity 1*-D* for *Campylobacter porAf2* variants identified amongst each of the flocks. A value of 1 indicates that all variants in a flock were different, and 0 indicates that all variants within a flock were identical.

The *porAf2* variant that became dominant in flocks 3, 14 and 19 was different in each of the flocks (Figures 1 and 2). These were *porAf2* variants 3, 8 and 736. The *porAf2* variant that became dominant in one of these flocks was present in the other two flocks at low frequency on two of the three occasions.

### Differences in Campylobacter porAf2 variant population structure between flocks, farms and parent flocks

With the exceptions of flocks 3, 14 and 19, the population structure was not distinguishable between individual flocks or farms. Flocks 3, 14 and 19 can been seen clustered to the extremes of the ordination plots, reflecting the dominance of different *porAf2* variants, but even then, they are not distinct from others (Figure 4a, b and c). Of the 245 *porAf2* variants identified amongst the study, 105 (42.9%) were detected on all three farms (Figure 4d). In total, they accounted for 93.2% of *porAf2* variants identified from samples from Farm 1, 92.1% of *porAf2* variants identified from Farm 3, and 79.9% of *porAf2* variants identified from Farm 2. Individually, the farms had between 8 and 39 unique *porAf2* variants, accounting for between 1 and 2.4% of the *porAf2* variants identified from each farm in total. A subset of broiler flocks were derived from matching parent flocks, and the *porAf2* variants were compared between parent-matched flocks for evidence of vertical transfer, either directly, or indirectly *via* faecally contaminated transport crates for example. In fact, between 1 (<0.1%) and 29 (2.2%) *porAf2* variants were unique to each group of broiler flocks with matched parents (Figure 4e) and parent-matched flocks could not be distinguished from each other using Bray-Curtis dissimilarity indices (Figure 4f).

**Figure 4.**
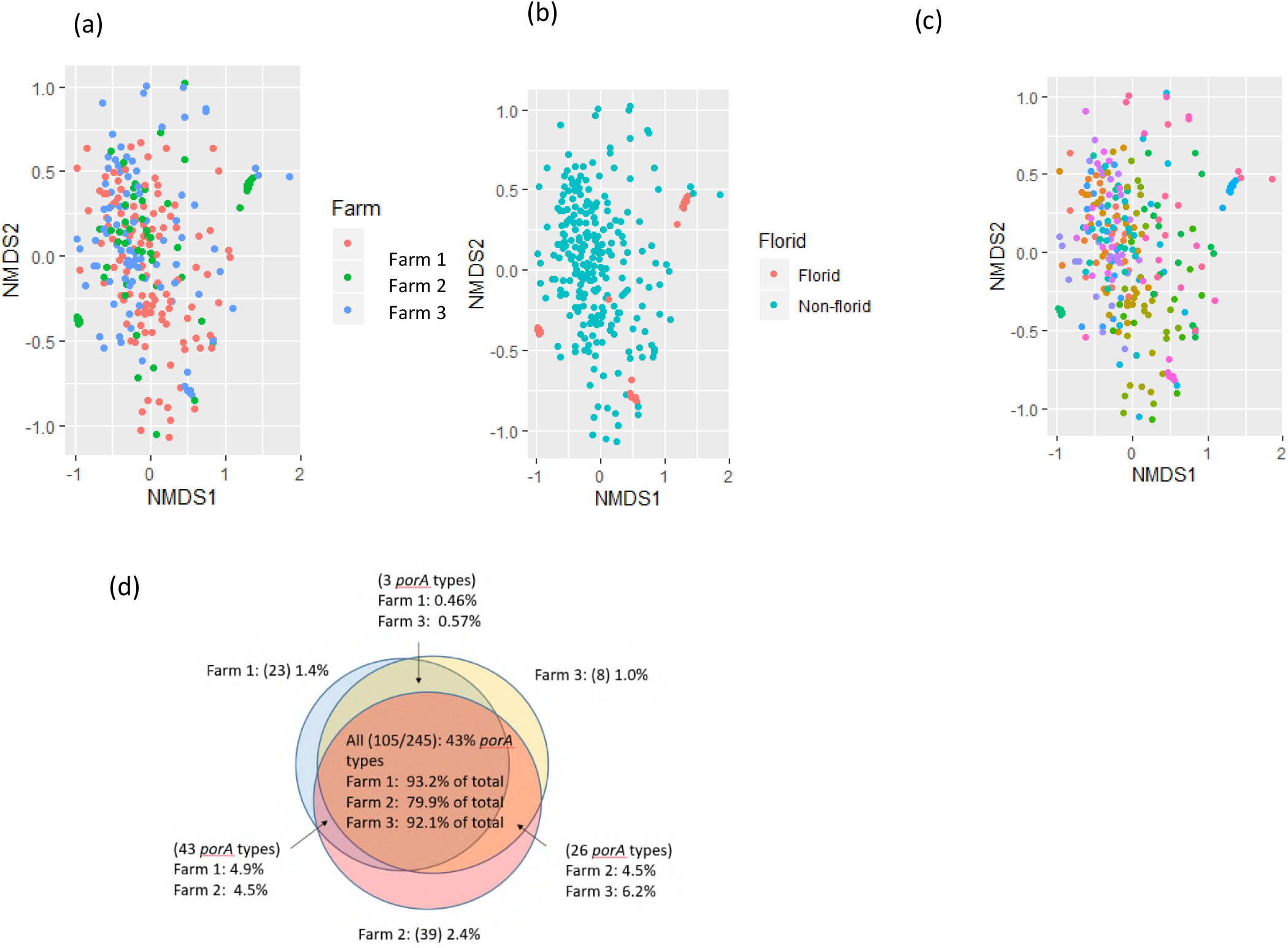
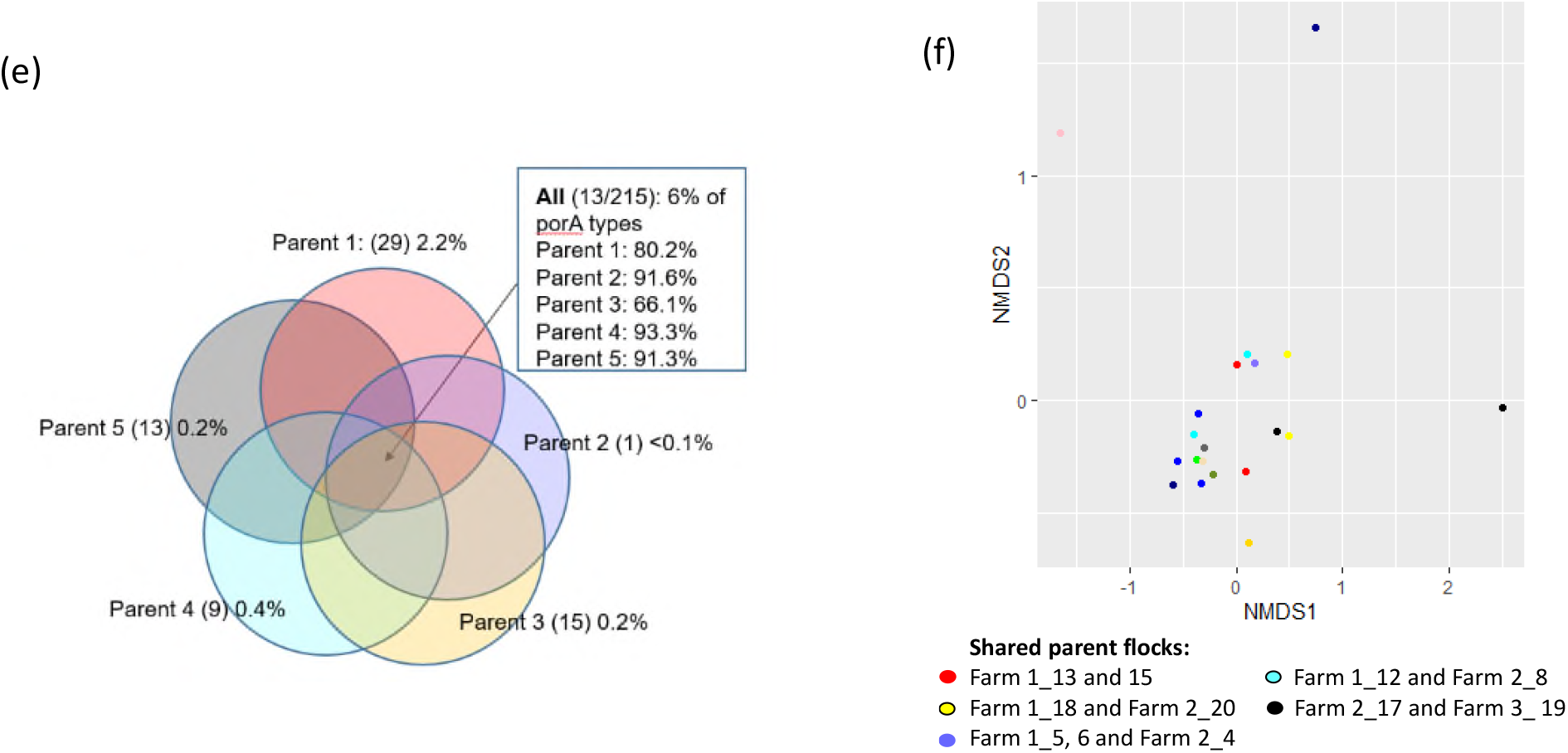
Non-metric multidimensional scaling (NMDS) ordination plots showing Bray-Curtis dissimilarity measures between the *porAf2* populations identified (a) farm, (b) ‘florid’ *versus* ‘non-florid’ status (c) flock (each different colour representing a different flock). Venn diagrams showing the overlap of *porAf2* variants identified by farm (d) and parent flock for a subset of 11 flocks (e). NMDS plot showing Bray-Curtis dissimilarity measures between the *porAf2* population detected amongst flocks derived from matched parent flocks (f).

## DISCUSSION

Results from this study show that *Campylobacter* DNA could be detected amongst faecal samples from 85-100% of the boiler flocks tested using nucleotide sequence based methods, depending on which gene target (*porAf2* or 16S) was used. The results were irrespective of whether or not the flocks may be called *‘Campylobacter* positive’ by routine culture or qPCR testing methods. Rather, the pattern of overgrowth of a predominant *porAf2* type coincided with flocks that would usually be determined *Campylobacter* qPCR/culture positive using Ct thresholds, whilst a pattern of high *porAf2* diversity at low prevalence was apparent amongst *Campylobacter* qPCR/culture negative flocks. The findings support the strong correlation between predominant *porA* type amongst culture/qPCR positive birds/flocks and diverse *porA* types amongst culture/qPCR negative birds/flocks shown in our previous work (Colles, et al., 2021). Two of the flocks had a small proportion of samples that were positive by conventional qPCR, but *Campylobacter* 16S nucleotide sequence was detected at very low frequency, if at all, with the *porAf2* pattern remaining diverse. Both of these flocks later tested positive for *Campylobacter* by standard culture methods in the abattoir, and may have been in early stages of transition when we tested them in this study. More work is needed to understand how a flock transitions from negative to positive for *Campylobacter* by conventional testing means, and could be critical for the control of *Campylobacter* infection in poultry flocks.

That *Campylobacter* DNA was not equally detected for samples tested by both 16S and *porA* is not surprising given they are different targets, with *porA* specifically amplifying *Campylobacter* DNA, and therefore giving a greater sensitivity of detection. In contrast, the 16S bacterial profile, amongst which *Campylobacter* in our studies represented <0.01% of species recovered from the broiler chicken faecal samples (Colles, et al., 2021) gives a high chance that more prevalent species will be detected ahead of *Campylobacter*, and may give greater variation between samples. Unlike our previous studies however, *Campylobacter* DNA was detected at a prevalence >1% of the bacterial profile in 14.8% (43/290) of samples in this study, and at 51-79% of the bacterial profile in 1.7% (5/290) of samples collected from the Swiss broiler flocks. Other studies report *Campylobacter* prevalence in excess of 10% for experimentally infected chickens (Han, et al., 2016; Rychlik, 2020). It is possible that some birds within the flocks were unusual in some way, for example with gut dysbiosis brought about by co-infection with another pathogen. Farm 3, with two *Campylobacter* qPCR/culture positive flocks from which the 5 samples with greatest *Campylobacter* prevalence were identified, was suspected to be having some problems with coccidial infections over the course of this study.

The *porAf2* variants 1-9 accounted for 70.5% of the total sequences recovered, and it was not possible to distinguish the *porAf2* populations isolated from broiler flocks by either farm or parent flock. Of the 245 *porAf2* variants identified amongst the study, 43% were identified from flocks on all three farms, with only 1-2.4% (8-39 *porAf2* variants) of the total, unique to each of the farms. These results imply that a large proportion of the *porAf2* variants are well-adapted for persistence in the poultry industry and co-exist amongst broiler flocks from different farms over a number of months. On a broader context, MLST-based studies also demonstrate a number of *Campylobacter* types show host association with chicken sources (Sheppard, et al., 2010). Deep sequencing using the *porAf2* target gives a useful indication of *Campylobacter* diversity within a sample, but further development of the method, including additional gene targets is required for more refined strain typing. It is not currently possible to distinguish between *C. jejuni* and *C. coli* species using the *porAf2* fragment, though the presence of both species amongst samples was detected by qPCR and it is likely the 80% homology between the most disparate nucleotide sequences reflects the different species also.

It was notable that nucleotide changes detected in the *porAf2* led to a new amino acid sequence (MOMPf2) in 70% of cases, with up to 79 *porAf2* nucleotide/65 MOMPf2 variants detected from an individual sample. This finding fits with the short variable region of the outer membrane protein being under host immune selection (Cody, et al., 2009), and the pattern of high diversity may reflect evasion of the host immune response (Bloomfield, et al., 2021). It was unpredictable which *porAf2* variant would become predominant in the three *Campylobacter* qPCR/culture positive flocks. Two of the *porAf2* variants colonising the *Campylobacter* qPCR/culture positive flocks were commonly observed amongst all flocks, implying they could have greater fitness, but the third was not. The stochastic nature of *Campylobacter* infection by different strain types has been shown to relate to the susceptibility of individual birds infected by a *Campylobacter* type by chance, more than the *Campylobacter* strain type itself, using Bayesian modelling approaches (Rawson, et al., 2020).

In conclusion, we demonstrate that *Campylobacter* testing methodology may need to be reviewed, depending on the circumstances in which it is needed. Whilst it is still useful to identify those flocks that are most heavily contaminated by *Campylobacter* at slaughter and pose greatest risk to human health by conventional means that are timely and cost-effective, identifying routes of transmission amongst a relatively rare gut inhabitant is much more challenging. The ability to routinely detect low levels of *Campylobacter* amongst broiler flocks at a much earlier age than would conventionally be identified requires a re-examination of how and when biosecurity measures are best applied. In addition, other interventions, such as good animal welfare and management to maintain optimal gut health may help to prevent the overgrowth of a single *Campylobacter* type, characteristic of problematic flocks that test *Campylobacter* positive by conventional means.

## ACKNOWLEDGEMENTS

This work was supported by the Biotechnology and Biological Science Research Council (grant numbers BB/N023803/1 and BB/K004468/1) including part of the Animal Health and Welfare ERA-net call. ALS was also supported by the UK Department for Environment, Food and Rural Affairs, (grant number OD0221). The collection of faeces was funded by the Federal Office of Food Safety and Veterinary Affaires (No. 2.16.03). We also acknowledge the support of the Bell AG, Zell, Switzerland, and the five farmers who provided access to their barns and logistic support.

